# Minimal Data · Maximal Insight (MDMI): A Structure-guided Pipeline for Discovering Functional Alternatives in Peptide-Protein Interfaces

**DOI:** 10.64898/2026.07.13.737974

**Authors:** Pouriya Bayat, Spencer J. Perkins, Sebastian Clancy, Sahil S. Patel, Richard F. Yin, Krištof Bozovičar, Serena Singh, Suman Shrestha, Zain Moustafa, Riham Zayani, Idorenyin IWE, Sepehr Bayat, Paul Kelly, Justin R.J. Vigar, Vivian Y. White, Matthew Xie, Mohammad Simchi, Sean Palter, Jessica Nguyen, Ilan Y. Zeisler, Bonny Wu, Keith Pardee

## Abstract

Discovering functional peptides across vast sequence space remains a formidable challenge, particularly when experimental training data is scarce. We present Minimal Data Maximal Insight (MDMI), a two-stage structure-guided computational pipeline that designs functional peptide variants using only a small, annotated dataset. Rather than relying on sequence information alone, MDMI integrates three-dimensional structural features derived from predicted peptide-protein complexes into a machine learning model that captures interface geometry and binding energetics. This structure-aware predictor, paired with a genetic algorithm for sequence exploration, reduced false positives from 70% to close to zero in an all-negative benchmark panel compared with a sequence-only model in computational benchmarking, and produced approximately four-fold more high-confidence *in silico* binders than state-of-the-art peptide/protein design baselines. Using the split-GFP system as a testbed, where fluorescence provides a direct functional readout of peptide-protein complementation, MDMI identified peptides with up to 38% sequence divergence from wild-type in Stage 1 while retaining measurable activity. In Stage 2, motif-guided recombination of successful Stage 1 variants produced highly divergent yet functional peptides bearing over 50% sequence difference from wild-type, revealing two distinct functional clusters in sequence space. As further validation, a top-performing candidate expressed as a full-length GFP fusion retained a GFP-like emission profile, supporting formation of a fluorescent GFP-like scaffold. These results demonstrate that structure-informed pipelines can uncover remote functional sequence space from minimal data, with broad implications for peptide and therapeutic analog discovery.

## Introduction

Peptides, short chains of typically 2 to 50 amino acids, participate in a wide range of biological processes, from endocrine signaling and neurotransmission to innate immunity and host defense^[1]^. Such peptides mediate a significant fraction of protein-protein interactions in cells (estimated up to ∼40%)^[2]^ and have gained prominence as therapeutics and research tools. Clinical interest in these molecules has surged, driven by their favorable pharmacology, enhanced tissue penetration, low immunogenicity as well as simpler solid-phase synthesis and the commercial success of peptide drugs such as GLP-1 receptor agonists^[3–8]^.

Despite this promise, systematically exploring the vast sequence space of peptides for new binders remains a formidable challenge. A peptide of moderate length (e.g., 20 amino acids) has on the order of 20^20^ possible permutations if only considering natural amino acids, an astronomical diversity that far outstrips what can be tested experimentally. Brute-force library screening methods (such as phage display or yeast display) can sample up to ∼10^9^-10^11^ variants in a single experiment, which is an infinitesimal fraction of all sequences^[9,10]^. In practice, peptide and protein libraries are often designed as focused or semi-rational libraries around known sequences or consensus motifs, since fully random sequence libraries contain only a very small fraction of folded or functional variants^[11,12]^.

Machine-learning (ML) frameworks such as RFpeptide^[13]^, Pepflow^[14]^, PepMLM^[15]^, and Peptune^[16]^ have been developed to narrow the search and accelerate design. Most are optimized primarily around binding, activity labels, or sequence plausibility, rather than around the precise bound geometry required for downstream functionality. Yet several practical constraints often limit ML-guided peptide design. First, most architectures demand thousands of labeled examples to generalize^[17,18]^. Attaining such large-scale experimentally measured properties (e.g., binding affinity, antimicrobial potency, or cell permeability) is rarely attainable for new peptide targets, especially outside of industry^[19]^. When only a few dozen such pairs exist, models tend to memorize rather than extrapolate^[20]^.

Moreover, unlike large proteins, peptides lack extensive structural databases to inform modeling. The Protein Data Bank contains hundreds of thousands of protein structures, but relatively few high-resolution peptide-protein complex structures^[21,22]^. This scarcity of structural data is problematic because short peptides typically do not adopt a fixed folded structure on their own; their functional conformations emerge upon binding to partner proteins^[23–25]^. Ignoring the context of the target interface can therefore mislead optimization, especially for flexible peptides that can bind in multiple conformations (so-called degenerate binding modes). Indeed, peptide activity is often highly context-dependent, and purely sequence-based design may stumble if a peptide’s binding modes or interface contacts are not explicitly considered^[26–28]^.

Finally, purely data-driven or deep generative models (e.g. large protein language models or diffusion-based protein designers) may struggle to propose truly diverse solutions: they tend to generate sequences similar or close to those seen in training or in natural proteins, and without careful tuning they might miss highly divergent yet functional peptides^[29–31]^. Overall, the dual problems of enormous sequence space and limited data/structural guidance have so far constrained our ability to discover diverse functional variants in peptide-protein interfaces.

In this study, we show that structure-aware modeling can uncover remote functional peptide variants even under low-data conditions. We focus on the challenge of identifying peptide analogs that preserve function while sharing as little as 50% sequence identity with the original peptide. To address this problem, we introduce Minimal Data · Maximal Insight (MDMI), a two-stage structure-guided design workflow that uses minimal experimental data to explore distant regions of the peptide sequence-function landscape.

As a testbed, we chose the split Green Fluorescent Protein (GFP) system, which is comprised of a 16-residue peptide (GFP_11_) and a larger GFP_1-10_ fragment, which only become fluorescent when bound together^[32]^ (Fig 1a.). The MDMI workflow is focused on the discovery of functional GFP_11_ analogs. This system provides a direct functional readout (fluorescence brightness) for peptide-protein binding and the interaction is well-defined and established in biotechnology^[32,33]^. In the first stage of our approach, we trained a machine learning model on a public GFP mutagenesis dataset^[34]^, from which we extracted approximately 100 variants whose substitutions were confined to the GFP_11_ segment and for which brightness measurements were available. Crucially, instead of relying only on sequence information, we incorporated structural features by predicting the 3D structure of each variant in complex with GFP_1-10_ using AlphaFold-Multimer^[35]^. From these predicted peptide-protein structures, we extracted interface descriptors including knowledge-based statistical potentials (from SPServer^[36]^) and physics-based energy terms (from PyRosetta^[37]^). These features capture complementary aspects of binding (evolutionary contact preferences and atomic interaction energies) and served as input to an ML model trained to predict GFP_11_ functionality (fluorescence).

**Figure 1.**
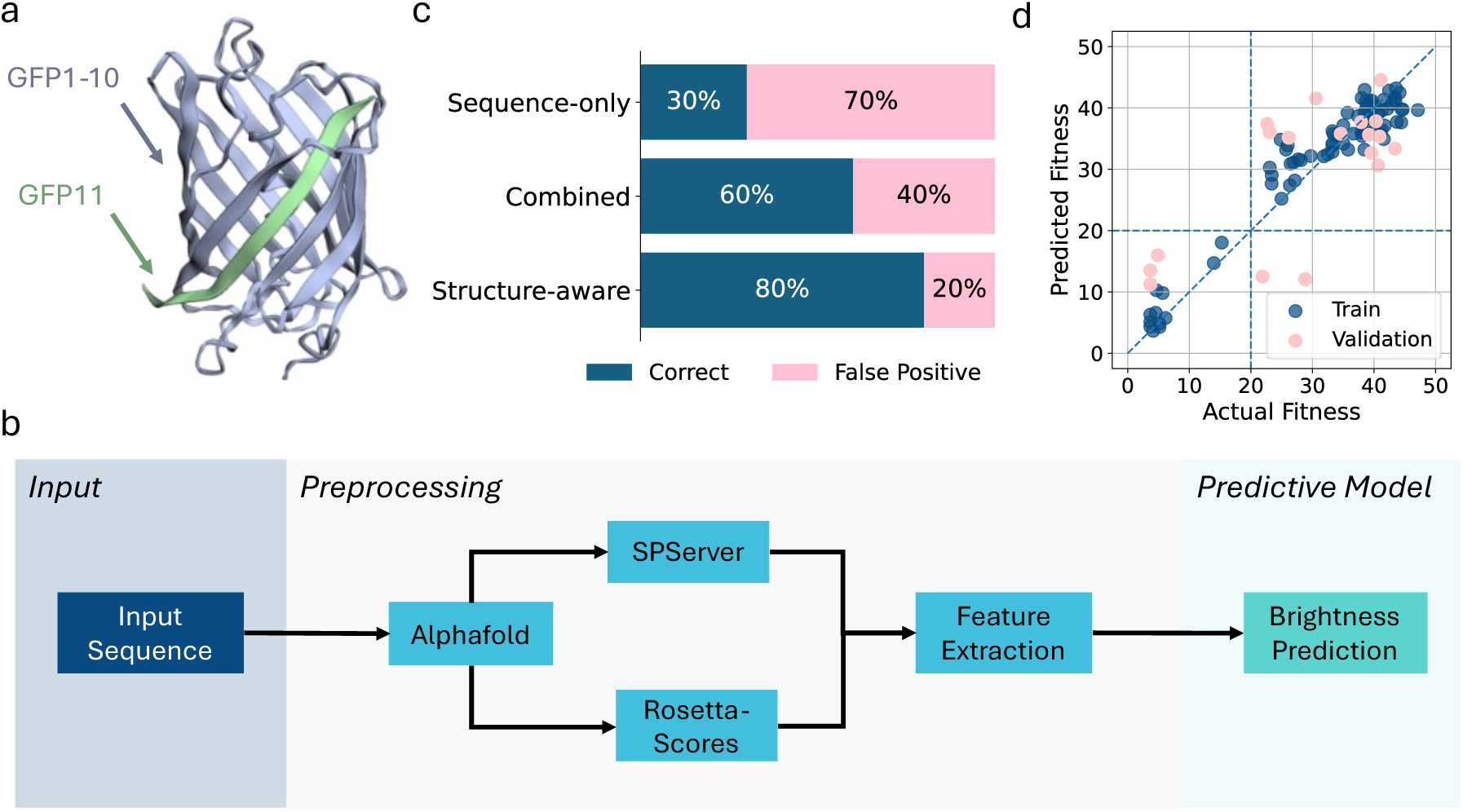
Overview of the MDMI (Minimal Data Maximal Insight) framework for peptide discovery. a. Structural model of the split-GFP system used in this study. The GFP11 peptide (green) binds to the larger GFP1-10 fragment (blue), and the degree of complementation correlates with fluorescence. b. Workflow for building the predictive model. Input peptide sequences are folded in the presence of GFP1-10 with AlphaFold-Multimer. Structural features are extracted using Rosetta scoring functions and SPServer metrics. These features are used to train a model predicting peptide-dependent fluorescence. c. Model specificity on an all-negative test panel. The sequence-only model misclassifies 70% of dim peptides with poor fluorescence while the structure-aware model reduces this false positive rate to 20%. d. Performance of the trained model. Predicted versus actual experimental fitness values are shown for both training (blue, train) and validation (pink, val) datasets.

Using this approach, even with the limited training examples, the structure-informed model more effectively rejected known non-binders than a sequence-only model, indicating that intermolecular geometry improves candidate classification under low-data conditions. We then used this predictive model to guide a genetic algorithm search through the sequence space. Starting from the wild-type GFP_11_ sequence, the algorithm iteratively generated new peptide variants (through mutations and recombination) and selected those predicted to maintain high fluorescence in the resulting GFP_11_/GFP_1-10_ complex. This yielded a set of moderately divergent but functional GFP_11_ candidates. We compared our model against RFdiffusion^[38]^, the current state-of-the-art zero-shot generator, as well as PepMLM^[15]^, a protein language model-based algorithm for structure-independent peptide binder generation, and observed that MDMI produced approximately four-fold more high-confidence binders as determined by their iPTM scores.

In the second stage of design, we analyzed the successful variants from the first stage to identify recurring mutational motifs (amino acid substitutions or patterns) that contributed to binding. By recombining these beneficial motifs, we constructed a next generation of highly mutated GFP_11_ sequences that were far more diverged from wild-type than any single-round variant. Strikingly, several of these designed peptides, containing well over 50% sequence difference from the original GFP_11_ peptide, retained robust fluorescence in experiments. Using this method, we obtained GFP_11_ variants bearing up to ∼63 % mutation relative to the wild-type while preserving function. While this phenomenon is observed in nature^[39,40]^, this outcome demonstrates that functional binding can be maintained within unique sequence space when computationally designed with informed constraints.

Our study illustrates a practical strategy for peptide engineering with a minimal training data set. By integrating *a priori* structural predictions into the learning and the search process, we exposed new regions of sequence space that brute-force or sequence-only approaches may otherwise miss. The resulting GFP_11_ variants show that short peptides can be highly evolvable *in silico* and able to diverge dramatically in sequence while preserving function, provided design algorithms account for the biophysical interaction requirements. We anticipate that this two-stage Minimal Data, Maximal Insight approach will be applicable to other systems where exhaustive screening is impractical and training data is limited, with application in the discovery of novel peptide and therapeutic analogs that remain functional even at a distance from known sequences.

## Results

We developed a two-stage strategy, Minimal Data · Maximal Insight (MDMI), to identify functional but highly divergent GFP_11_ variants. In Stage 1, a structure-aware machine-learning model trained on a minimal public dataset guided a genetic algorithm toward functional sequence space, generating candidates with 3 to 6 mutations predicted to retain fluorescence. The top candidates were synthesized and screened experimentally. In Stage 2, recurring beneficial mutations from the validated Stage 1 hits were recombined to construct new peptides. This yielded variants with >50% sequence divergence from wild-type GFP_11_ that still complemented GFP_1-10_ with strong fluorescence, revealing multiple distinct sequence solutions for preserving function.

### Stage 1: Structure-Aware Modeling Enables Enrichment of Functional Variants

We hypothesized that explicit integration of structural interaction data could overcome the data scarcity limitations for peptide binder design. To test this, we first drew on a publicly available Aequorea victoria GFP (*av*GFP) mutant dataset providing brightness measurements^[34]^. From this resource, we filtered 109 variants bearing substitutions exclusively in the GFP_11_ segment (aa 215 - aa 230). Our workflow comprised two main components: a predictive model and a generative model (Fig. 1b-c).

### Structure-Aware Predictive Modeling

The predictive arm of the workflow begins with 3D structural simulation using AlphaFold-Multimer^[35]^, generating peptide-protein complexes for the GFP_11_ variants. In our initial model iteration, we analyzed these complexes using SPServer alone, which provides knowledge-based statistical potentials that assess interatomic interactions^[36]^. As the model underwent multiple revisions (Supplementary Table S1.), we evolved from using a public dataset with SPServer-only scoring to incorporating our own experimental dataset and adding PyRosetta physics-based energy terms such as van der Waals, solvation and hydrogen-bond terms^[37]^. Next, we fed the extracted structural descriptors into a machine-learning pipeline designed to learn low-dimensional, functionally relevant embeddings (Fig. 1b). After preprocessing, the selected SPServer- and PyRosetta-derived features were encoded with a variational autoencoder (VAE) composed of fully connected layers with batch normalization and SELU activations. A downstream fully connected neural network (FCNN) then used these latent representations to predict peptide fluorescence upon complementation with GFP_1-10_.

We first trained a regressor on continuous fluorescence intensities (labels) derived solely from the structural features and then applied a fluorescence threshold separating functional and non-functional peptides only at evaluation time. This two-step paradigm is widely used in quantitative-structure-activity-relationship (QSAR) modeling, where models are trained on continuous activity values and a biologically defined activity threshold is applied afterward to classify compounds as active or inactive^[41,42]^. In practice, the regressor produces a continuous score, and the activity cutoff acts as a post-hoc classification threshold. To validate that structural features provide essential discrimination capability beyond sequence information alone, we tested model specificity using non-binding peptides against a sequence-based model. This test reveals the model’s tendency to incorrectly classify non-binders as positive hits, a critical concern in drug discovery where false positives waste experimental resources. The sequence-only model exhibited a 70% false positive rate (7/10 negatives misclassified), whereas the structure-aware model (SPServer only) achieved 20% false positive rate (2/10 negatives misclassified), demonstrating that structural features capture critical binding determinants that sequence patterns cannot reliably identify. Adding an ensemble layer that combined an XGBoost regressor trained on the original structural features with a KNN regressor operating on VAE-derived latent embeddings eliminated all false positives (Fig. 1d). On the training and hold-out sequences the best performing ensemble achieved: Root Mean Square Error (RMSE) = 3.57 (train) / 9.36 (val); accuracy = 100%/88.9%; Matthews correlation coefficient (MCC) = 1.00 / 0.75; Area Under the Receiver Operating Characteristic Curve (AUROC) = 0.98 / 0.91; Area Under the Precision-Recall Curve (AUPRC) = 0.99 / 0.98 (Supplementary Table S2). The training coefficient of determination was R^2^=0.92, while the validation performance dropped to R^2^=0.49, indicating imperfect quantitative calibration. However, the rank ordering of sequences was preserved, as reflected by AUROC = 0.91. Because downstream decision-making depends only on whether a candidate exceeds the fluorescence threshold, classification metrics (Accuracy = 88.9%, MCC = 0.75) are the most relevant indicators of performance.

### Genetic Algorithm for Sequence Generation

With the goal of creating peptides that have limited similarity to the original (wild-type) peptide, in parallel we implemented a genetic algorithm via the PyGAD framework^[43]^. The genetic algorithm served to explore the broader sequence space systematically, drawing on our predictive model to guide which productive mutations to retain (Fig. 2). Beginning with a randomized population of peptide sequences, each peptide was evaluated for its predicted brightness and complex stability with GFP_1-10_. Sequences with greater fitness underwent crossover and mutation operations, thereby creating offspring peptide sequences that inherit advantageous structural and functional traits. By iterating through several generations, the genetic algorithm converged on top-ranked variants, which were then prioritized for empirical validation. During selection, predicted solubility was assessed using the CamSol^[44]^ web server, and only candidates with both favorable solubility scores and high predicted brightness were retained for wet-lab testing.

**Figure 2.**
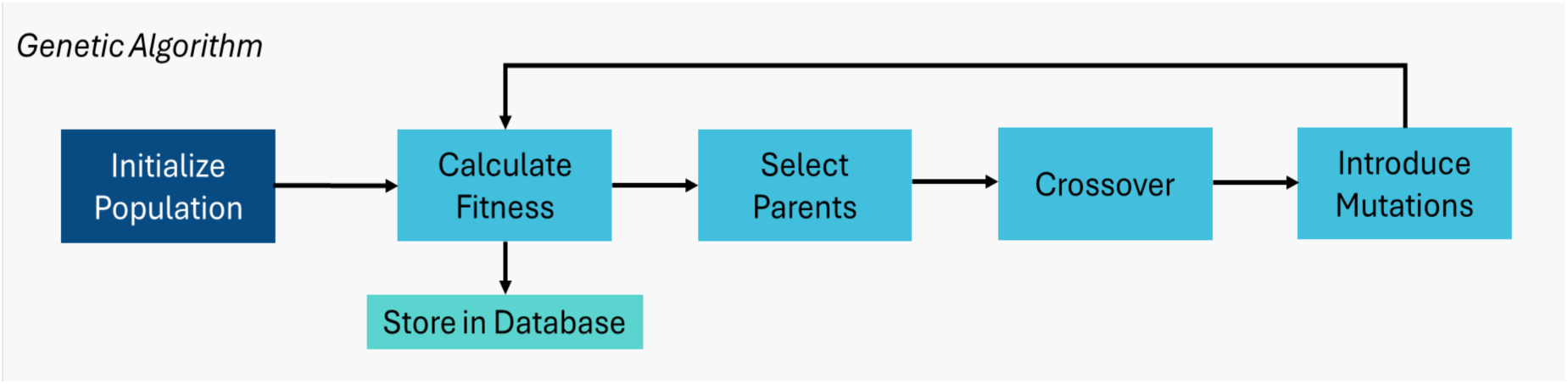
Overview of the genetic algorithm for sequence diversification. A genetic algorithm from PyGAD initializes a population of candidate sequences, evaluates their predicted fitness using the model, and iteratively improves them through selection, crossover, and mutation.

### Retraining Our Model Using In-House Data

After we established confidence in the predictive model using a public dataset (Fig. 1d)^[34]^, we then used the model to generate new candidate sequences, which were subsequently evaluated by in-house functional screening across a range of fluorescence outputs (Supplementary Fig. S1). Building on this, we generated our own experimental dataset to retrain the model and implemented a genetic algorithm to design new sequences. The original dataset, sourced from the literature^[34]^, involved *in vivo* expression in *E. coli* of *av*GFP mutants. We hypothesized that a dataset collected under conditions consistent with our *in vitro* experiments would enhance the model’s performance.

Consequently, we screened 114 GFP_11_ variants comprising sequences generated by the initial model trained on the public dataset, together with randomly mutated control sequences. Their fluorescence was measured relative to wild-type GFP_11_ after complementation with purified superfolder GFP_1-10_ in a cell-free expression assay, generating the in-house dataset used for retraining.

The screening results and position-wise amino-acid distributions for the 114-sequence in-house retraining set are shown in Fig. 3a-b. Figure 3c reports the positional distribution of substitutions across the entire screened library. Substitutions were concentrated at positions 7, 9, and 11 (110/114, 113/114, and 109/114 variants, respectively), while the other mutable positions were represented at lower but broadly similar frequencies (16 to 59 variants per site). The glutamate residue (E8) in the center of the peptide plays an important role in coordinating the chromophore responsible for fluorescence and was held invariant by design^[45,46]^. Given the role of the E8 region in chromophore formation, variation at adjacent residues is expected to affect fluorescence in both positive and negative ways. Although the wild-type GFP_11_ is moderately polar (GRAVY score of -0.10)^[47]^, our mutant peptides span the entire spectrum: some retain a balanced profile (51/114), while others skewed slightly toward hydrophobic (47/114) or hydrophilic (16/114) content (Fig. 3d).

**Figure 3.**
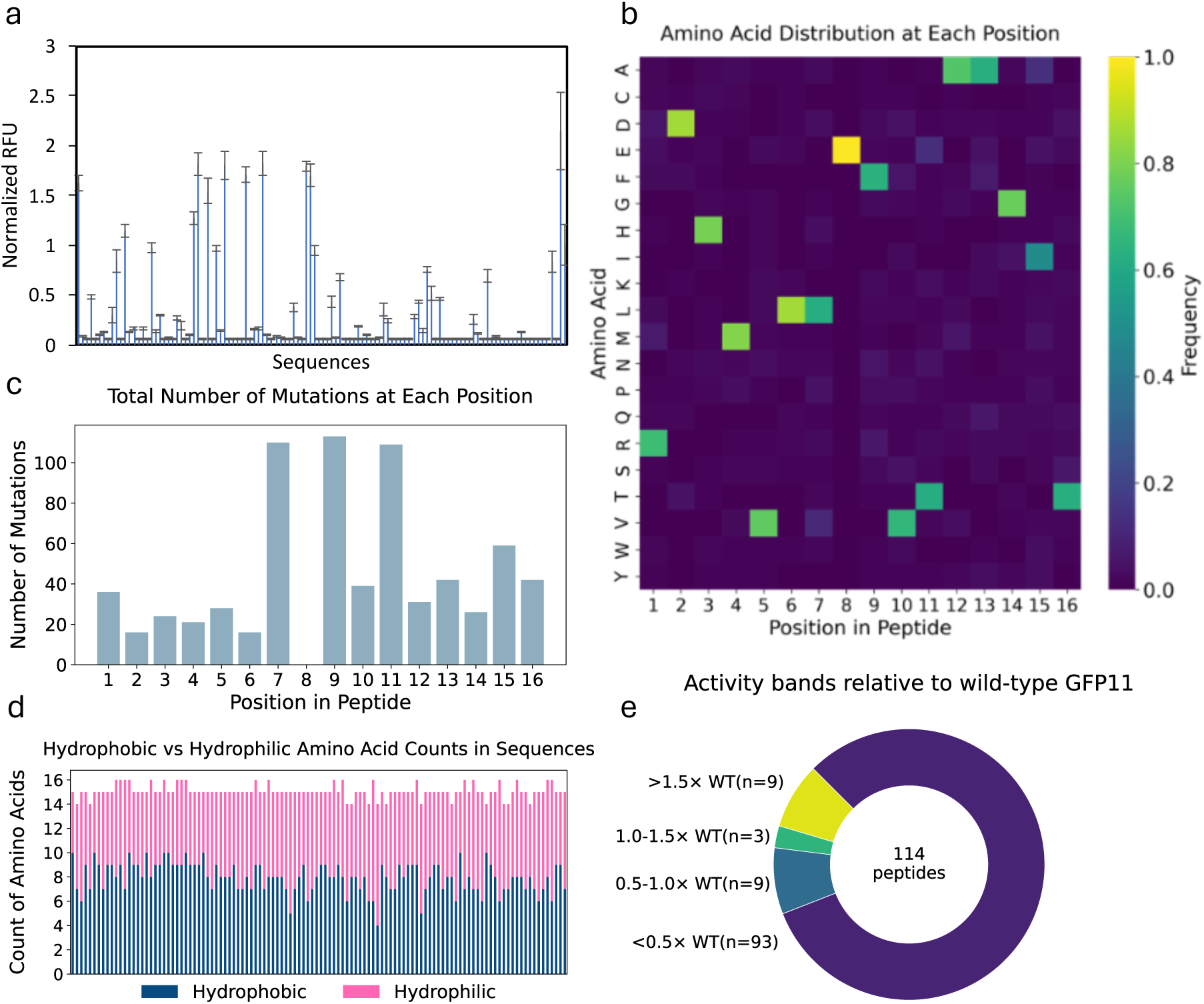
Characterization of the in-house experimental dataset used to retrain the predictive model. a. Normalized fluorescence (RFU) values for peptides synthesized and screened in-house. These measurements were used to retrain the model on experimentally relevant functional data. Wild-type (WT) is equal to 1. Bars represent mean ± standard deviation, n=3 b. Heatmap showing amino acid frequencies at each position across the peptide sequences in the dataset, highlighting positional variability. c. Histogram of mutation counts relative to the wild-type GFP11 sequence. d. Hydrophobic vs. hydrophilic amino acid content per sequence, showing a broad distribution of physicochemical properties in the dataset. e. Distribution of peptide variants across four fluorescence activity bands (<0.5×, 0.5-1.0×, 1.0-1.5×, and >1.5× wild-type brightness). Most variants fall below 0.5× wild-type brightness, whereas a small subset matches or exceeds wild-type levels.

Most peptide variants were substantially dimmer than wild-type GFP_11_, although a small subset matched or exceeded wild-type brightness. As a result, the retraining set contained examples spanning strongly deleterious to strongly enhancing sequence changes, allowing the model to learn from graded fluorescence outcomes rather than binary labels. At the same time, the strong skew toward weak variants meant that the retraining set was dominated by low-activity examples, leaving relatively sparse coverage of the highest-performing sequences.

For visualization, fluorescence values were expressed as fold brightness relative to wild-type GFP_11_ and grouped into four activity bands: <0.5×, 0.5-1.0×, 1.0-1.5×, and >1.5× wild-type brightness (Fig. 3e). Most variants were substantially dimmer than wild-type, with 93 of 114 peptides (81.6%) falling in the lowest activity band, whereas 12 of 114 peptides (10.5%) matched or exceeded wild-type brightness. This distribution highlights the scarcity of high-performing variants in the retraining dataset, while still providing both strongly deleterious and markedly enhancing examples for model learning.

### Benchmarking MDMI against RFdiffusion and PepMLM

Using the newly generated dataset (Fig. 3), we retrained our model and ran the genetic algorithm to find another generation of novel functional variants (Fig. 4). We benchmarked the sequences generated by MDMI using the metrics reported by AlphaFold in comparison to RFdiffusion, a general-purpose zero-shot backbone generator conditioned on target structure^[38]^, as well as PepMLM, a sequence-only span-masked language model trained on 10000 protein-peptide binders^[15]^.

**Figure 4.**
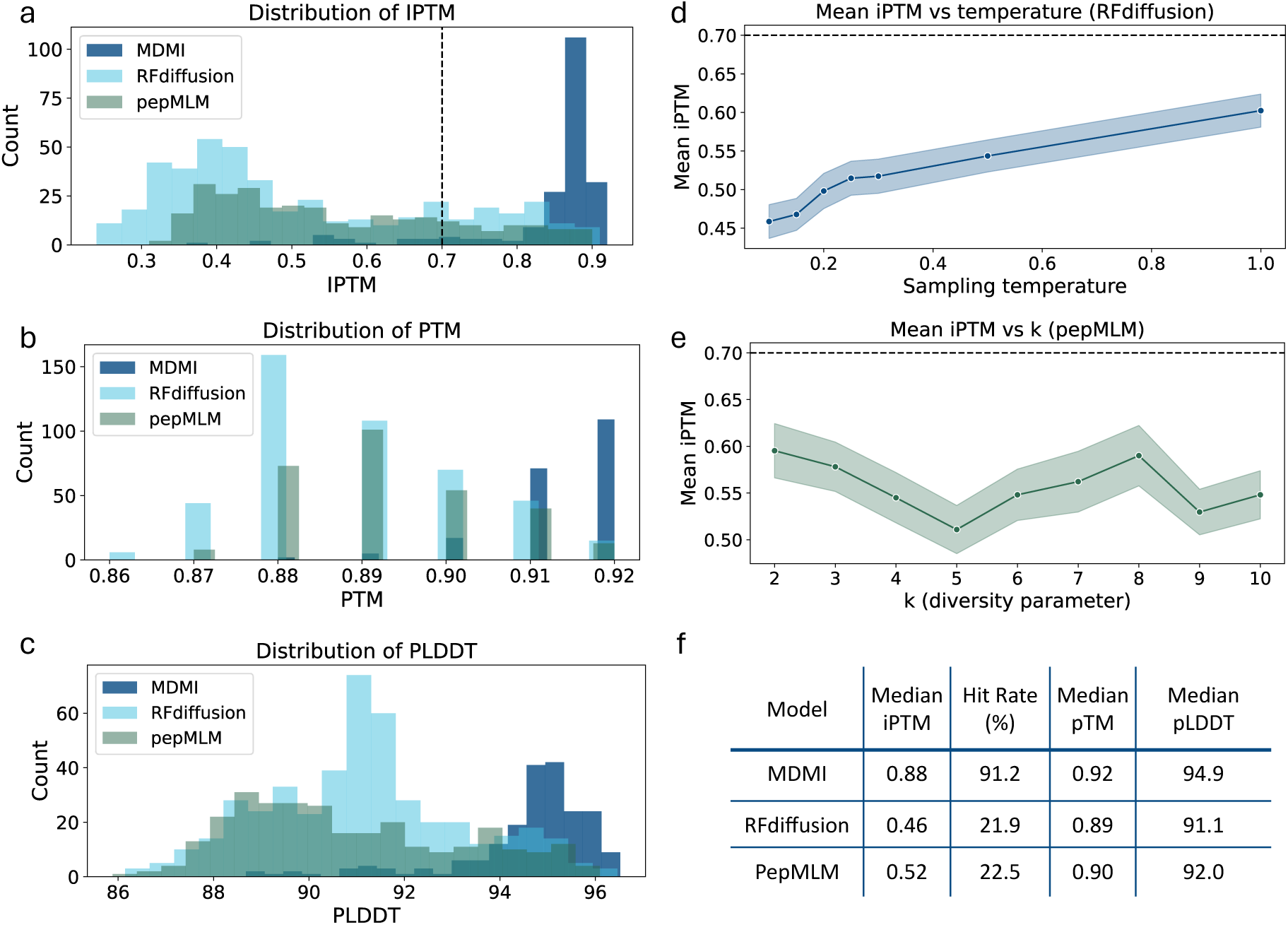
Computational quality assessment. a. Distribution of interface TM-scores (iPTM) predicted by AlphaFold-Multimer for MDMI (navy), RFdiffusion (cyan), and pepMLM (forest green) designs. The dashed line marks the *in silico* success threshold (0.7). b. Predicted TM-score (pTM) distributions for the same design sets. c. Per-residue backbone confidence (pLDDT) distributions. d. RFdiffusion designs generated at increasing sampling temperatures show a modest upward trend in mean iPTM. Shaded band indicates ±1 standard error of the mean (SEM); the dashed line marks the *in silico* hit threshold (iPTM = 0.7). e. PepMLM designs sampled with top-k = 2-10 show a non-monotonic relationship between diversity and quality. Mean iPTM dips at k = 5, recovers above k = 6, and peaks at k = 8. Shaded band: ±1 SEM. f. Summary statistics for each method. Table reports the median iPTM, hit rate (percentage of designs with iPTM ≥ 0.7), median pTM, and median pLDDT.

For benchmarking, GFP1-10 was held fixed as the receptor sequence for all methods, and each method was asked to generate 16-residue peptide candidates matching the length of GFP_11_. Because RFdiffusion and PepMLM were evaluated across parameter sweeps rather than as one-candidate-per-target generators, the total number of sampled candidates differed between methods (MDMI, n = 204; RFdiffusion, n = 448; PepMLM, n = 289). To avoid post hoc cherry-picking, we retained all generated candidates and evaluated every design using the same AlphaFold-Multimer confidence metrics namely, Template Modeling (TM)-score metrics (iPTM and pTM) and residue-level confidence (pLDDT); pTM, for the accuracy of the model’s predicted topology for GFP_11 variant/_GFP_1-10_ overall, iPTM, for gauging how reliably the model positions the interacting subunits relative to one another, an indicator of interface credibility and potential binding affinity^[15,35]^, and pLDDT for residue-level structural integrity (Fig. 4a-c, f).

In this *in silico* benchmark, the MDMI output set (n = 204) showed a median iPTM of 0.88 (inter-quartile range [IQR] 0.86-0.90), whereas the RFdiffusion set (n = 448) had a median of 0.46 (IQR 0.37–0.63) and PepMLM (n = 289) reached 0.52. Thus, both baseline methods produced substantially lower-confidence interfaces than MDMI overall. Applying an *in silico* threshold of iPTM > 0.7 for effective binding (i.e., the predicted interface between peptide and target is well-formed and likely to be physically realistic)^[48,49]^, 91.2% of MDMI designs qualified as “hits” compared with 21.9% for RFdiffusion and 22.5% for PepMLM.

In other words, MDMI produced roughly four-fold more high-confidence binders than either baseline. This advantage was also reflected in backbone-confidence metrics: MDMI achieved a median pTM of 0.92 and median pLDDT of 94.9, compared with 0.89/91.1 for RFdiffusion and 0.90 / 92.0 for PepMLM (Fig. 4b-c, f). Together, these results indicate that MDMI produces peptide-GFP_1-10_ complexes with higher-confidence binding interfaces and stronger overall backbone confidence than either baseline, yielding peptide-GFP_1-10_ complexes that are more likely to adopt plausible bound conformations.

Because RFdiffusion is a stochastic generator, its output quality can depend on the sampling temperature (T), which controls the degree of randomness during structure generation with higher values increasing diversity. To assess whether MDMI’s advantage depended on a particular RFdiffusion setting, we generated peptide-GFP_1-10_ complexes across discrete sampling temperatures from 0.10 to 1.00 (Fig. 4d). Across this range, iPTM showed only a modest positive correlation with temperature (Pearson r = 0.24), suggesting that this approach may slightly benefit the design process. Designs sampled at T = 0.20 yielded 18.8% hits (12/64), whereas those at T = 1.00 yielded 34.4% hits (22/64). Thus, although higher temperatures modestly improved RFdiffusion output within the tested range, even the best-performing setting remained substantially below the hit rate achieved by MDMI.

We performed an analogous analysis for PepMLM to determine whether its benchmark performance depended strongly on the sequence-sampling regime. In PepMLM, top-k sampling restricts each masked position to the k most probable amino acids, thereby controlling the tradeoff between determinism and sequence diversity. Across top-k = 2 to 10, interface quality varied non-monotonically (Fig. 4e): mean iPTM decreased at intermediate values and then rebounded, peaking at k = 8. Excluding the single deterministic sequence generated at k = 1, hit rates ranged from 15% to 34%, with k = 8 producing the best yield (34%). These results indicate that PepMLM performance is somewhat sensitive to sampling strategy, but that reasonable variation in top-k does not close the gap to MDMI.

### Experimental validation of the model trained with in-house data

Using the in-house retraining dataset and the updated structure-aware predictor, and seeking to develop GFP_11_ peptides that are unique from wild-type sequences and validated experimentally, we implemented the genetic algorithm to identify sequences with 3-6 mutations, corresponding to approximately 19-38% mutation rates. With the design goal focused on peptide-protein interaction, the catalytically essential glutamate (E8) has been held constant. Across 1327 *in silico* variants explored by the genetic algorithm (Fig. 5a-c), the distribution of model-predicted fluorescence exhibits a steadily rising upper tail, with the 95th percentile predicted fluorescence increasing by ∼39% from the earliest to the latest candidates, indicating that although the bulk of the population hovers near the starting baseline, successive rounds of selection and mutation uncover and preserve peptides with higher fluorescence scoring (Fig. 5a). The accompanying amino-acid frequency heat-map (Fig. 5b) reveals enrichment at select positions (e.g., 1, 2, 11, and 15) where a subset of residues appears more frequently, suggesting selective pressure at these sites while other positions tolerate diverse substitutions, indicating localized flexibility within the binding interface.

**Figure 5.**
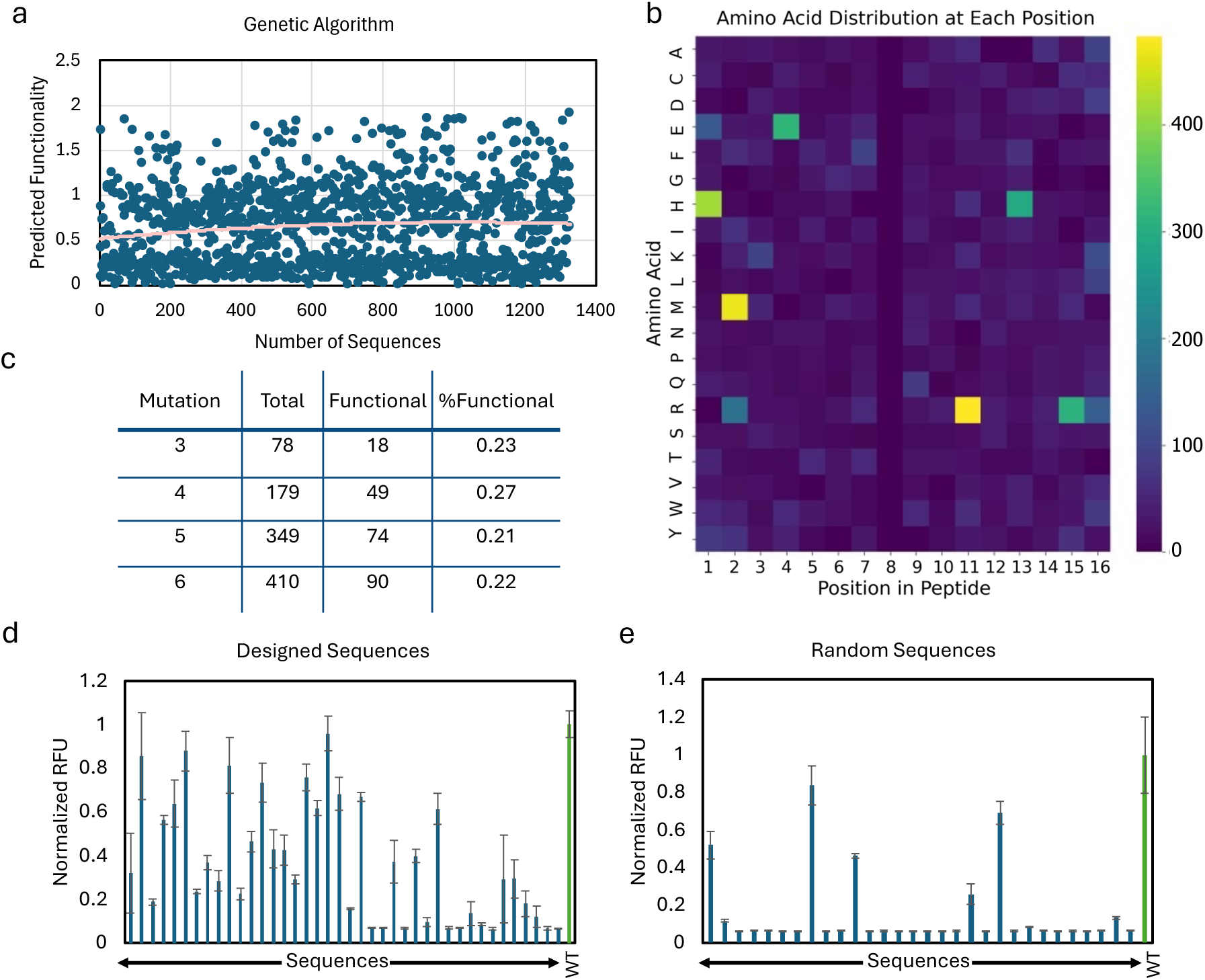
Summary of the Stage 1 genetic algorithm guided designs. a. Predicted functionality of sequences over generations in the genetic algorithm. While most sequences remain near baseline, the model identifies variants with consistently higher predicted functionality. b. Heatmap showing amino acid frequency across all generated sequences, highlighting convergence at specific positions (e.g., 1, 2, 11, 15) despite diverse overall sequence composition. c. Distribution of generated sequences by mutation count, with corresponding counts of predicted functional variants and their relative proportions. Functionality rates remain consistent across different mutational depths. d. Experimental fluorescence (normalized RFU) of designed peptides (without pLM conditioning). A range of brightness values is observed, with several sequences approaching wild-type levels. e. Experimental fluorescence of randomly mutated sequences assayed during construction of the in-house retraining dataset, shown as a baseline reference for the fluorescence distribution obtained by unguided mutagenesis, highlighting the non-triviality of functional peptide discovery and the relative enrichment achieved by the design process (Green bar indicates the Wild-type (WT); blue bars, test peptides - Bars represent mean ± standard deviation, n=3).

From the pool of functional variants with 3-6 mutations, we selected the top 40 sequences with the highest predicted fluorescence and estimated solubility scores for experimental validation. These variants were expressed using linear DNA templates in a cell-free protein expression system to assess their functionality when mixed with purified GFP_1-10_. As illustrated in Figure 5d-e, the sequences within the designed batch exhibited significantly higher signals than the randomly mutated peptides sampled during construction of the retraining dataset. Specifically, the area under the curve (AUC) for the designed batch was 2.7 times larger for the designed sequences. Furthermore, 63% of the designed sequences exhibited more than 20% activity relative to the wild-type GFP_11_ peptide, 28% showed over 60% GFP_11_ activity, and 20% demonstrated greater than 75% GFP_11_ activity compared to the wild-type. Together, these results show that retraining on the in-house dataset and coupling the model to the genetic algorithm enriched for functional GFP_11_ variants.

### Integration of a Protein Language Model

State-of-the-art peptide design often incorporates protein language models (pLMs) to capture sequence-level evolutionary features that may aid mutagenesis and function prediction. To test whether such information improves MDMI, we added a ProtBERT-based branch to the workflow, using the Rostlab/prot_bert_bfd checkpoint and an evo-tuning strategy inspired by Biswas et al.^[50]^ in which the model was first adapted on fluorescent-protein family sequences and then fine-tuned on experimentally measured GFP_11_ functionality (Fig. 6a). Embeddings from this branch were compressed and combined with the structural feature pathway to enable direct comparison with the previously introduced non-pLM model.

**Figure 6.**
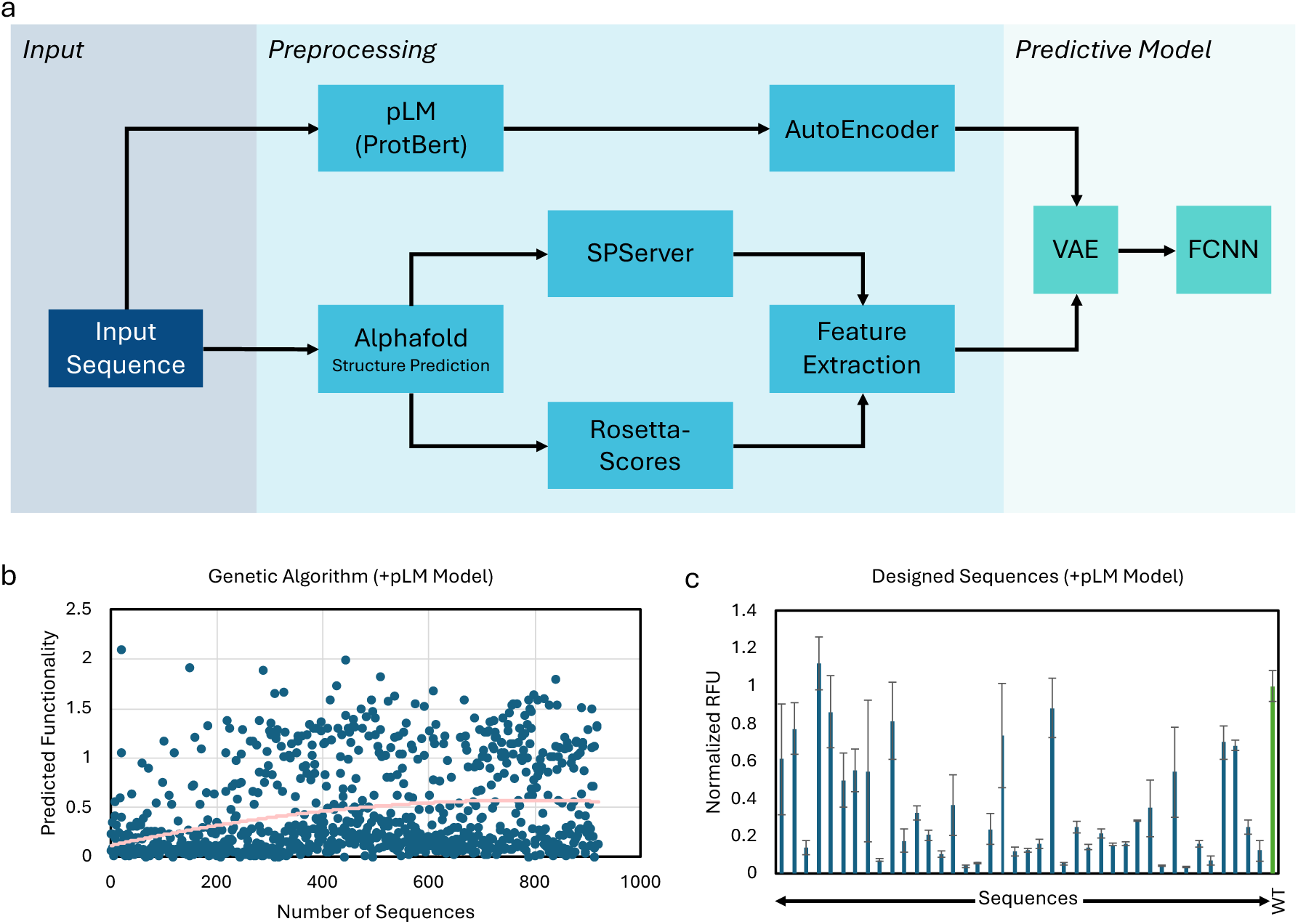
pLM-guided designs and screening. a. Schematic workflow of the predictive part of MDMI equipped with a protein language model (pLM) b. Predicted fitness values for sequences generated by the genetic algorithm using the pLM-guided model. The red trend line indicates a modest upward trend over generations, reflecting steady exploration of functional space. c. Experimental fluorescence (normalized RFU) of pLM-guided designed peptides. Functional outcomes were broadly similar to non-pLM designs, but sequences were identified more rapidly, suggesting improved search efficiency rather than enhanced fitness (Green bar indicates the Wild-type (WT); blue bars, test peptides - Bars represent mean ± standard deviation, n=3).

To begin, the pLM-equipped model was paired with the genetic algorithm to generate novel GFP_11_ variants in the same 3 to 6 mutation regime used previously (approximately 19 to 38% sequence divergence). Across the *in silico* search, the 95th percentile of model-predicted fluorescence increased by ∼52% over the genetic algorithm trajectory, indicating progressive enrichment of higher-scoring candidates (Fig. 6b). We then selected 40 sequences with high predicted functionality and favorable solubility for experimental validation and screened them using cell-free expression from DNA templates encoding the variant peptides.

Experimentally, however, the pLM-guided set shifted toward weaker outcomes overall than the non-pLM set (Fig. 6c). The fraction of peptides exceeding 20% of wild-type GFP_11_ activity decreased from 63% to 56%, while the proportions reaching the higher activity thresholds remained broadly similar to those obtained without the pLM. In other words, adding the pLM increased the number of low-activity candidates without improving the recovery of the strongest performers. Together, these results suggest that, in the current low-N regime, the structurally informed MDMI workflow does not benefit from incorporating a pLM.

### Stage 2: Motif-Guided Assembly of Highly Divergent Variants

Encouraged by Stage 1’s ability to find functional peptides with up to 38% divergence (3-6 mutations; Fig. 5-6), we next decided to test a higher degree (>50%) of sequence divergence from wild-type GFP_11_. Such highly mutated sequences are attractive for discovering alternative binding modes, orthogonal peptide tools, or flexibility around intellectual property but they represent a remote sequence space where machine learning predictions are often unreliable. This challenge was particularly stringent because the predictor had been trained predominantly on peptide variants with 5 to 7 mutations from wild-type GFP_11_ (median = 6 mutations, 37.5% divergence; IQR = 6 to 7 mutations), with only sparse coverage of more highly divergent variants (8 to 9 mutations) and these highly divergent variants were uniformly weak in the assay. Consistent with this limitation, when we directly asked the model to propose GFP_11_ sequences with >50% mutations (i.e. 8 or more substitutions), the resulting designs uniformly failed to reconstitute fluorescence (Supplementary Fig. S3).

These results highlight a practical limit of purely algorithmic design: once candidate sequences move well beyond the mutational regime represented in the training data, prediction becomes an extrapolation problem and both false positives and false negatives become more likely. Similar generalization challenges have been seen in other ML-guided protein design studies, where models can extrapolate modestly beyond their training regime but show a marked drop in functional yield when pushed deeper into sequence space^[51]^. To overcome this hurdle, we returned to leverage motif-guided recombination of the best Stage 1 variants. We hypothesized that, while our single model-driven attempt to jump to >50% sequence divergence was not successful, the clues about how to build a functional but distantly related peptide may be contained within the designs of successful moderate GFP_11_ mutants. By analyzing the top-performing variants, defined here as those retaining >60% of wild-type fluorescence,

we identified recurring amino acid substitutions that could serve as transferable design motifs. A heatmap of these bright binders revealed several positions with non-wild-type residues enriched at high frequency (Supplementary Fig. S4). Guided by these patterns, we manually assembled a new set of GFP_11_ sequences, each carrying a combination of the most enriched substitutions. In total, we designed ten highly divergent peptides, each containing 10 mutations relative to wild-type GFP_11_ (i.e. only 6 of 16 positions retained, ∼63% sequence difference).

The 10 variants along with the wild-type and a non-binder peptide were chemically synthesized and complemented with purified GFP_1-10_ at final concentrations of 71 µM (peptide) and 106 µM (GFP_1-10_) in PBS (total volume 16 µL; molar ratio ∼1.5:1, GFP_1-10_:peptide). Notably, nine of the ten highly divergent peptides generated measurable fluorescence, and several reached activities close to wild-type (Fig. 7a, Supplementary file) indicating that extensive mutation does not necessarily compromise function and demonstrated that the split-GFP interface tolerates significant sequence diversity.

**Figure 7.**
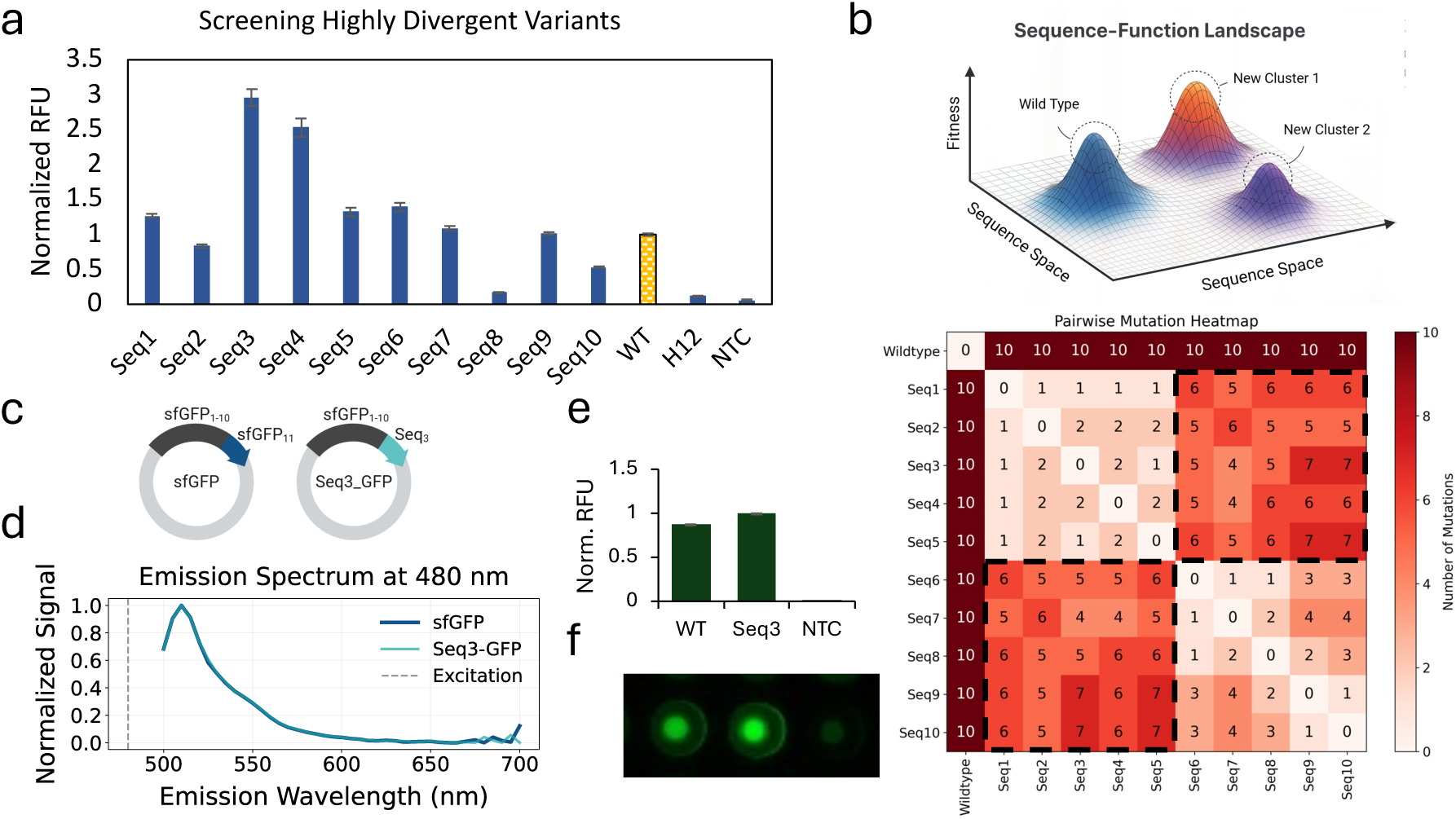
Stage 2 motif-guided designs. a. Normalized fluorescence (relative fluorescence units, RFU) obtained when the variants were mixed with purified GFP1-10. Wild-type (WT, yellow), non-binder (H12), and no-template control (NTC) are shown for reference b. Sequence-level organization of the highly divergent GFP11 variants. Top, conceptual schematic illustrating the emergence of distinct functional sequence clusters in sequence space. Bottom, pairwise substitution matrix for Seq1-Seq10 and wild-type GFP11, showing two internally related but distinct sequence families. c. Schematic of the full-length constructs used for follow-up analysis. In the Seq3-sfGFP construct, the wild-type GFP11-derived eleventh-strand segment was replaced with the highly divergent Seq3 sequence. d. Emission spectra of purified WT (sfGFP) and Seq3-GFP, showing closely similar spectral profiles. The dashed line indicates the excitation wavelength. e. Normalized fluorescence of purified WT (sfGFP) and Seq3-GFP. f. Fluorescence image of purified proteins dispensed in a 384-well plate. WT (sfGFP) and Seq3-GFP show strong fluorescence, whereas the negative control remains dim.

To explore how far we had travelled through sequence space, we mapped the ten variants onto a pairwise substitution matrix (Fig. 7b). Every variant carries ten substitutions relative to wild-type GFP_11_ (10/16 residues), yet the matrix reveals discrete clusters. Seq1-Seq5 form a tightly knit group that differ from one another at ≤ 2 positions, whereas Seq6-Seq10 constitute a second group, again internally similar, but separated from the first by 4-7 substitutions. Thus, motif-guided assembly did not merely recover a single local solution, but produced multiple functional and distant sequence families.

As an exploratory follow-up, we asked whether one representative highly divergent sequence could also support fluorescence in the context of the full-length sfGFP scaffold. We constructed two expression plasmids: a canonical sfGFP control (GFP_1-10_ + GFP_11_) and a fusion construct, Seq3-GFP, in which the Seq_3_ sequence was appended to the C-terminus of GFP_1-10_ in place of the wild-type GFP_11_ (Fig. 7c). Both constructs were expressed in *E. coli* BL21(DE3) and purified by His-tag affinity chromatography. Spectral analysis showed that Seq3-GFP retained an emission profile closely matching that of sfGFP, indicating that the Seq3 variant supports formation of a GFP-like fluorescent scaffold (Fig. 7d; full spectra in Fig. S5). Endpoint fluorescence measurements at equivalent protein concentrations further confirmed that Seq3-GFP produced strong fluorescence, comparable to, or slightly exceeding the wild-type sfGFP (Fig. 7e-f). Notably, this comparable performance in the intramolecular fusion context is consistent with the expectation that the potential advantage observed in the bimolecular complementation assay (Fig. 7a) becomes reduced when peptide-protein proximity is enforced by covalent linkage.

## Discussion

Data scarcity has long hampered predictive modeling of proteins and peptides because the combinatorial explosion of admissible sequences quickly outpaces feasible wet-lab throughput. Prior approaches either relied on pretrained language models that assume the target sequence distribution resembles natural proteins, or on large, supervised datasets that are generally unattainable for newly emergent targets. In contrast, MDMI exploits a structure-aware model from a few dozen labeled examples. The success we observe with the split-GFP system suggests that high-quality, interaction-aware features can compensate for limited sample size; this principle will likely translate to other peptide-protein interfaces provided that the complex can be modeled with reasonable confidence. In benchmarking against state-of-the-art models (e.g. RFdiffusion and PepMLM), MDMI produced approximately four-fold more high-quality binder sequences that exceeded an iPTM threshold of 0.7. This improvement is notable because all three methods were evaluated using the same AlphaFold-Multimer confidence metrics. The higher hit rate therefore reflects the fact that MDMI proposes a candidate pool that is already enriched for sequences compatible with a high-confidence GFP_1-10_ interface, whereas RFdiffusion and PepMLM generate broader sets of candidates that contain many more low-confidence designs.

Our results demonstrate that MDMI can navigate peptide sequence space effectively when only ∼100 experimentally annotated variants are available. By employing features derived from protein-peptide complex scoring, the workflow retained sufficient sensitivity to rank-order GFP_11_ mutants (AUROC = 0.91, Fig. 1d) and to steer a genetic algorithm that yielded functional variants (Fig. 5). In the second stage of the workflow, motif-guided recombination of successful Stage 1 designs produced functional peptides with >50% sequence divergence from wild-type GFP_11_ (Fig. 7). This two-stage strategy therefore expanded the search beyond the local mutational neighborhood while avoiding overreliance on direct extrapolation. The highly mutated sequences were manually assembled by combining residues that were repeatedly enriched among the top hits, which currently limits throughput and leaves room for user bias. This requirement for human intervention likely reflects the fact that the model could identify local sequence-function patterns within the sampled regime but could not reliably extrapolate them into remote sequence space without an additional motif-level constraint. Future versions of MDMI could automate this step with crossover or fragment-shuffle operators so that exploration of distant sequence space is fully algorithmic.

Although MDMI succeeds in a controlled split-GFP setting, several caveats merit attention. First, the workflow inherits the uncertainty of single-structure prediction: AlphaFold-Multimer provides a single dominant pose for each peptide-target pair, but peptide binding in solution can involve conformational heterogeneity, side-chain repacking, and local induced fit. Consequently, some candidates may appear favorable because of one high-scoring static pose, even though that interaction is not stable across an ensemble of conformations. This issue is especially important for highly mutated peptides, where binding may depend on subtle induced-fit effects. Future versions of MDMI could address this by incorporating features derived from docking and molecular dynamics.

Second, the computational cost of structure prediction remains non-trivial (∼5 min per sequence on a high-end GPU, e.g., NVIDIA® RTX™ A6000). A more scalable next step would be to use multi-stage screening, in which inexpensive surrogate models or sequence-level filters triage large candidate pools and AlphaFold-Multimer is reserved for a smaller, higher-value subset.

Third, looking toward the future with the incorporation of non-natural amino acids for therapeutic peptide design, our experiments focus on canonical amino acids; extending MDMI to non-natural residues will likely require explicit parameterization of those residues in the underlying modeling stack, including new residue templates, rotamer libraries, and score-function or charge terms, or alternatively the use of learned physicochemical-property surrogates derived from chemical structure^[52–54]^. Enabling non-natural residues would substantially broaden the accessible design space and could allow MDMI to optimize not only binding, but also properties such as proteolytic stability, bioavailability, and target selectivity. Finally, although the genetic algorithm achieved substantial enrichment, alternative search strategies such as Bayesian optimization and reinforcement learning have also been applied to protein and peptide sequence design and may further accelerate convergence in larger sequence spaces^[55–57].^

High-affinity peptide binders underlie modalities ranging from biosensors to targeted drug delivery to therapeutics. By lowering the data threshold for productive modeling, MDMI can streamline discovery programs where scant experimental data is the chief constraint. In practice, we envision coupling MDMI with iterative rounds of cell-free protein expression-based screening to realize a closed-loop design-build-test cycle. Because the design framework is modular, it could easily be adapted to new target peptide-protein pairs. In such applications, only the target-specific scoring function would need to be updated for the new peptide target, and the machine learning model could be retrained using the small number of hits found through directed evolution campaigns, such as phage or mRNA display experiments^[58,59]^.

In summary, MDMI shows that by using 3D structural modeling along with a machine learning model, we can find functional peptide sequences even when very little data is available. As platforms for low-volume screening mature, such structure-aware pipelines may become increasingly useful for rational peptide engineering.

## Methods

### Predictive Model

The predictive framework comprises a preprocessing pipeline and a two-branch neural network that integrates structural, statistical, and sequence-based features to predict fluorescence output for novel GFP_11_ variants.

### Data Preprocessing

The dataset was partitioned pseudo-randomly into training (80%), validation (10%), and test (10%) sets using scikit-learn’s train_test_split utility. To maximize training data for model learning, the validation and test subsets were drawn from the original 20% holdout. Structural features were generated via Local ColabFold, a ColabFold implementation of AlphaFold2 utilizing MMseqs2 for multiple sequence alignment. GFP_11_ sequences were concatenated to GFP_1-10_ with a chain break indicator and processed in batches. Resulting PDB structures were scored using SPServer to extract 12 global normalized statistical potentials that quantify interfacial quality and penetration metrics. Additional energy terms and solvent-accessible surface area (SASA) metrics were computed using PyRosetta’s scoring functions. In total, 49 structural and statistical features were compiled.

To reduce model complexity, we performed feature importance analysis using ensemble approaches including SMOTE with random forest, gradient boosting, and XGBoost with custom loss weighting. The top 14 features, six from SPServer (Es3dc, E3dc, E3d, ZEcomb, ZEloca, ZE3dc) and eight from PyRosetta (fa_intra_sol_xover4, fa_rep, hbond_lr_bb, pro_close, average_energy, std_energy, total_energy, separated_sasa), were selected. All features were standardized to zero mean and unit variance using scikit-learn’s StandardScaler fitted on the training set and applied to validation and test sets.

### Variational Autoencoder (VAE)

A variational autoencoder was implemented in PyTorch Lightning to learn a compressed representation of the features. The encoder and decoder comprised alternating linear layers, one-dimensional batch normalization, and SELU activations, with a latent bottleneck of 16 dimensions. The encoder outputs parameterized mean (μ) and log-variance (log σ²) vectors, from which latent samples were drawn via the reparameterization trick. The VAE loss function combined mean squared reconstruction error, Kullback-Leibler divergence, and two regularization terms: a distance loss encouraging alignment between normalized latent distances and label-derived distances, and a line loss preventing linear correlation between latents and inputs. The relative weight of the distance term was increased progressively during training to promote informative latent representations. Optimization employed Adam (learning rate=0.01, weight decay=1e-5) with a learning rate scheduler (γ=0.993). Early stopping was applied with patience of 100 epochs and a minimum δ of 0.01; training was limited to 1,000 epochs. The best-performing model on the validation set was retained.

### Fully Connected Neural Network (FCNN)

Encoded latent vectors (16 dimensions) were input to a fully connected neural network for regression. The network, also in PyTorch Lightning, comprised linear layers of sizes 32, 64, 128, and 256, interleaved with batch normalization, SELU activations, and a dropout layer (p=0.2) after the first hidden layer. A final sigmoid activation scaled outputs to match the fluorescence range. Training utilized Adam (learning rate=0.002, weight decay=1e-6) with a scheduler (γ=0.98) and early stopping (patience=30, δ=0.01), for up to 100 epochs.

### pLM-Based Branch

A pre-trained protein language model was fine-tuned with a custom fine-tuning step as a way to provide information about evolutionary fitness. Based on work by Biswas *et al.*^[50]^, ProtBERT was trained using “evo-tuning”, using sequences from the fluorescent protein family to learn the distinct features of our target protein GFP_11_. Afterwards, a regular fine-tuning approach was applied to train a top-level model with experimentally determined data about GFP_11_ functionality. After training, generated protein embeddings from the last hidden layer outputs with a CLS pooling scheme were used to help guide the full predictor model. To make sure the weight of these features did not overwhelm the results from the structurally informed models, we trained a small auto-encoder model to compress the embeddings down to the twelve most important features, which were combined with the structural features in the VAE.

### Genetic Algorithm

To explore sequence space and optimize GFP_11_ fluorescence, we implemented a genetic algorithm using PyGAD^[43]^. An initial population of 20 randomly generated 16-mer sequences was encoded via the VAE and evaluated by the FCNN. After each generation, mutation statistics were updated in a dictionary mapping each residue substitution to a fitness weight. New sequences were generated by mutating the top 10 performers, using weighted probabilities that balanced exploratory random mutations and exploitation of successful substitutions. Mutation probabilities transitioned from uniform to dictionary-informed as the weight factor increased over generations. In the final optimization phase, mutation donors were selected from a tree structure of all prior generations. Each generation node scored by (mean top-ten fitness), success rate, and branching factor determined a propensity score. This approach preserved promising lineages while avoiding stagnation in low-fitness branches.

### Implementation and Computational Resources

All genetic algorithm computations and network inferences were executed on a workstation (32 cores, 256 GB RAM, NVIDIA RTX A6000). Sequence preprocessing and model evaluation required ∼5 minutes per sequence. A parallel work-pool architecture on Google Cloud VMs, coordinated via MongoDB, enabled near-linear scaling across 20 worker nodes.

### Expression and Purification of GFP1-10

The pET15b+ plasmid containing the GFP_1-10_ gene was ordered from Twist Bioscience and used to transform BL21(DE3) *E. coli* (New England Biolabs). A single colony was picked and grown overnight in 25 mL LB medium supplemented with ampicillin. A 1:100 dilution was used to inoculate 2 × 1 L LB medium containing ampicillin, and the culture was incubated at 37 °C until reaching an OD600 of ∼0.7. Expression was induced with 1 mM IPTG, and cultures were further incubated at 30 °C for 5 hours. Cells were harvested by centrifugation (4000 × g, 10 min, 4 °C), and the resulting pellet was stored at −80 °C until purification. The cell pellet was resuspended in buffer A (20 mM HEPES pH 7.6, 300 mM NaCl, 10% glycerol, and 1 mM DTT/0.5 mM TCEP) at a ratio of 10 mL buffer per gram of pellet. Cells were sonicated on ice (5 s ON, 25 s OFF, 40% amplitude, ∼1800 J for ∼45 mL lysate) and centrifuged (maximum speed, 2 × 30 min, 4 °C) to remove cell debris. Purification was performed using an ÄKTA Pure L system (Cytiva). The lysate was supplemented with imidazole to a final concentration of 10 mM before loading onto a pre-equilibrated HisTrap HP (Cytiva) column. The column was washed sequentially with buffer A + 20 mM imidazole and then 50 mM imidazole until reaching a baseline absorbance. Bound protein was eluted with buffer B (20 mM HEPES pH 7.6, 300 mM NaCl, 10% glycerol, 250 mM imidazole, 1 mM DTT/0.5 mM TCEP), pooled, and concentrated. The protein was buffer-exchanged into storage buffer (50 mM HEPES-KOH pH 7.6, 100 mM KCl, 10 mM MgCl₂, 20% glycerol). The protein was snap-frozen in liquid nitrogen and stored at −80 °C.

### Expression, Purification, and Characterization of sfGFP and Seq3_GFP Fusion Protein

To construct Seq3_GFP, the coding sequence of Seq3 was appended to the 3′ end of the GFP_1-10_ open reading frame and cloned into a pET BLANK (Amp) plasmid (Twist Bioscience). The canonical sfGFP construct, in which the wild-type GFP_11_ sequence was appended to GFP1-10 in the same vector backbone, was prepared in parallel as a control. Both constructs were expressed and purified following the same protocol described above for GFP_1-10_. Briefly, constructs were transformed into BL21(DE3) *E. coli* (New England Biolabs), expressed under IPTG induction, and the His-tagged fusion proteins were purified by Ni-NTA affinity chromatography. Protein concentration was determined by absorbance measurement at 280 nm using a NanoDrop spectrophotometer (Thermo Fisher Scientific), and samples were normalized to equal concentration prior to fluorescence measurements.

For plate-based fluorescence quantification, equal amounts of purified sfGFP and Seq_3__GFP were dispensed into a 384-well black-walled plate and fluorescence intensity was measured with excitation at 488 nm and emission at 530 nm. For spectral analysis, emission scans were collected from 450 to 700 nm at 350-540 nm excitation, and each spectrum was normalized to its peak value. For fluorescence imaging (Fig. 7f), wells containing purified proteins were imaged under blue-light illumination.

### Oligonucleotide Design and Assembly

Variant coding sequences for GFP_11_ (16 amino-acid peptide) were ordered as two complementary 60-nt single-stranded DNA oligos (IDT) that carry a T7 promoter, ribosome-binding site (RBS) and ∼10-nt reciprocal overlaps (Supplementary Fig. S2). DNAs were either ordered pre-resuspended or were resuspended after delivery in nuclease-free water (Fisher Scientific) to 100 µM, then combined 1:1 to give a 50 µM duplex-ready mix for each variant. PCR extension was performed afterwards to convert the oligos into a full double stranded oligo. PCRs (25 µL) contained 12.5 µL 2× Q5 Master Mix (New England Biolabs), 2.5 µL oligo mix (50 µM each strand), and 10 µL water. Cycling: 95 °C 2 min; 10-30 cycles of 95 °C 5 s, 57 °C 30 s, 72 °C 60 s; final extension 72 °C 4 min; 4 °C hold.

### Cell-free Expression Screening

Reactions were performed in a 384-well flat-bottom polypropylene plate (Corning 3544). A 15 µL master-mix was prepared on ice for each round as follows:

**Table 1.** Cell-free expression reaction components.

| Component | Volume μL (per reaction) |
| --- | --- |
| Solution A (NEB PURExpress®) | 6 |
| Solution B (NEB PURExpress®) | 4.5 |
| Murine RNase inhibitor (NEB) | 0.3 |
| Purified GFP<sub>1-10</sub> (8.2 μM) | 1 |
| Nuclease-free H<sub>2</sub>O | to 15 |

Immediately before dispensing, 1 µL of freshly assembled dsDNA template was added to the master mix, mixed, and the reaction was transferred to the plate. Controls on every plate included the wild-type as the positive control and a non-template sample as the negative control. Plates were sealed with optically clear film and incubated at 37 °C inside a plate reader. Fluorescence was recorded for 2-8 h using both bottom and top optics, excitation 488 nm, emission 530 nm, gain 75-100 (auto-range off).

## Acknowledgements

We would like to thank Dr. Yohei Yokobayashi and Dr. Samuel Hauf for their generous support during Pouriya Bayat’s research internship at Okinawa Institute of Science and Technology (OIST) and for stimulating discussions on peptide screening approaches that informed the early directions of this work.

